# Identifying signals of memory from observations of animal movements

**DOI:** 10.1101/2023.08.15.553411

**Authors:** Dongmin Kim, Peter R Thompson, David Wolfson, Jerod Merkle, L. G. R. Oliveira-Santos, James D. Forester, Tal Avgar, Mark A. Lewis, John Fieberg

## Abstract

Incorporating memory (i.e., some notion of familiarity or experience with the landscape) into models of animal movement is a rising challenge in the field of movement ecology. The recent proliferation of new methods offers new opportunities to understand how memory influences movement. However, there are no clear guidelines for practitioners wishing to parameterize the effects of memory on moving animals. We review approaches for incorporating memory into Step-Selection Analyses (SSAs), a frequently used movement modeling framework. Memory-informed SSAs can be constructed by including spatial-temporal covariates (or maps) that define some aspect of familiarity (e.g., whether, how often, or how long ago the animal visited different spatial locations) derived from long-term telemetry data. We demonstrate how various familiarity covariates can be included in SSAs using a series of coded examples in which we fit models to wildlife tracking data from a wide range of taxa. We discuss how these different approaches can be used to address questions related to whether and how animals use information from past experiences to inform their future movements. We also highlight challenges and decisions that the user must make when applying these methods to their tracking data. By reviewing different approaches and providing code templates for their implementation, we hope to inspire practitioners to investigate further the importance of memory in animal movements using wildlife tracking data.

## 1. Introduction

Animal movement impacts ecological processes at all levels, including individual foraging efficiencies (Bonte et al., 2012; Van Moorter et al., 2009), population persistence (del Mar Delgado et al., 2018; Miles et al., 2020), species distributions (Fortin et al., 2005; Matthysen, 2005), connectivity (Doherty & Driscoll, 2018) and ecosystem functioning (Bauer & Hoye, 2014; Nathan et al., 2008; Subalusky et al., 2017). While much research has explored how animal movements are influenced by environmental conditions (Fortin et al., 2005), intra- and inter-specific social interactions (Matthysen, 2005), and internal states such as hunger levels (Hooten et al., 2019), the importance of past experience and memory is also increasingly recognized as a key component of animal movement (Piper, 2011; Fagan et al., 2013; Lewis et al., 2021). For example, by remembering the location and outcomes of previously visited locations, many species can increase energy intake rates (Janson & Byrne, 2007; Trapanese et al., 2019) and avoid areas that might increase mortality risk (Bracis et al., 2018; Gehr et al., 2020; Heathcote et al., 2023). Further, by remembering average environmental conditions, such as the average timing of resource waves, animals can better time migratory movements (Abrahms et al., 2019; Bracis & Mueller, 2017). Thus, models that incorporate memory are important for both developing and testing ecological theory, and they are likely to lead to improved predictions of how animals will respond to changes in their environment (Fagan et al., 2023; Gurarie & Avgar. 2024).

One straightforward approach to integrating such complex types of memories into a model is to assume that past experiences can be encoded into a spatially-referenced system in the animal’s brain (sometimes referred to as a “cognitive map”), which is then accessed during the retrieval phase to inform movements. Although hidden from direct observation, a spatially referenced map can be mathematically modeled as a surface changing dynamically over time as memories are lost, reinforced, or replaced. These constructs are central to key empirical models for memory, including “time since last visit” to a location (Schlägel & Lewis, 2014) as a determinant of wolf movement (Schlägel et al., 2017), and episodic returns of brown bears to ephemeral seasonal resources (Thompson, Derocher, et al., 2022; Thompson, Lewis, et al., 2022). Overall, the map is a latent, spatially-referenced variable, whose dynamics are inferred indirectly from animal movement patterns.

The information that animals gather, and perhaps memorize, as they move can be divided into three categories: 1) spatial information (i.e., locations animals have visited), 2) site attributes, including resource quality or quantity, and 3) temporal information (i.e., about how long-ago animals visited a previous site or when a site peaks in forage quality), with #2 and #3 comprising ‘attribute memory’ (Fagan et al., 2013; Kashetsky et al., 2021; Lewis et al., 2021; Thompson, Lewis, et al., 2022; Tulving & Craik, 2000). Food-caching blue jays (*Cyanocitta cristata*) use all three kinds of information: they remember the locations of many caches, the type of seed in each cache, and how long it has been since making the cache (Darley-Hill & Johnson, 1981). Owing to the temporal variability present in most environments, it can be advantageous to rely more heavily on recent experiences and to discount memories from long ago (Tello-Ramos et al., 2019). For example, roe deer (*Capreolus capreolus*) primarily base their foraging decisions on recent experiences due to rapid changes in resource availability within their home ranges (Ranc et al., 2021). Note, however, that memory is also expected to temporally decay due to the limitations of the neurological infrastructure that holds it, and distinguishing such decay from an adaptive discounting may be particularly challenging. By revisiting sites, animals can update their knowledge of site attributes; optimal return times may depend on how quickly the reliability of past information decays due to environmental change as well as resource renewal rates (Ranc et al., 2021; Spencer, 2012). For example, wolves (*Canis lupus*) and brown bears (*Ursus arctos*) delay returning to previously visited kill sites so that prey numbers may recover (Gurarie et al., 2022; Selva et al., 2017).

Ecologists have developed theoretical models to explore how past experiences and memory might influence animal movements (Fagan et al., 2013; Forrest et al., 2024; Kashetsky et al., 2021; Lewis et al., 2021; Wang & Salmaniw, 2023). The simplest encodable memory attribute is familiarity with a given location, either whether an individual has ever visited the site (Northrup et al., 2016; Rheault et al., 2021) or how frequently it has visited the site in the past (Bracis et al., 2018). When coupled to spatial movement models, preference for familiar locations is sufficient for the formation of stable home ranges (Gautestad et al., 2013; Heathcote et al. 2023; Merkle et al., 2017; Nabe-Nielsen et al. 2013, 20; Ranc et al., 2022; Van Moorter et al., 2009). More complex memory attributes include locations of resources or past conflicts, allowing animals to integrate spatial and attribute memory (i.e., memory of where positive and negative experiences occurred). Attraction to previously discovered resources can lead to resource-driven patterns of nonterritorial spatial segregation (Aarts et al., 2021; Riotte-Lambert et al., 2015). By way of contrast, memory and avoidance of locations where past conflicts with conspecifics occurred can give rise to spontaneous territorial pattern formation (Potts & Lewis, 2016; Ellison et al., 2020).

Advances in animal tracking technology and statistical modeling approaches have motivated ecologists to explore the potential for memory-informed movements in a wide range of animal taxa (although terrestrial mammals, mainly ungulates, remain by far the most-studied group; see Table 1). By tracking individual animals over consecutive years, ecologists can identify whether current movements can be explained from previously observed locations.

**Table 1.** Empirical modeling for memory.

| Paper | Species | Memory representation | Types | Methods | Model fit | Habitat-selection predictors |
| --- | --- | --- | --- | --- | --- | --- |
| Avgar et al. 2015 | Woodland caribou | Dynamics in animals' habitat selection over landscape and time | Forage | Likelihood of animals moving with cognitive selection functions | Bayesian (MCMC) | 1) Foraging qualities<br>2) Predator density (Wolf)<br>3) Moose habitat<br>4) Snow depth |
| Rheault et al. 2021 | Mule deer | Differences between animals' occurrences throughout time | Forage | SSF framework with an OD as a response variable in the model | Step Selection Functions (SSFs) with conditional logistic regression | 1) Terrain ruggedness index<br>2) elevation<br>3) distance to treed edges<br>4) snow depth<br>5) land covers<br>6) NDVI<br>7) distance to well pads<br>8) distance to roads<br>9) previous ODs (past yr)<br>10) current ODs (prev 7 days) |
| Falcon-Cortes et al. 2021 | Elk | Number of visits to the foraging path | Forage | A Markovian model with the probability of moving to patches | Bayesian (MCMC) | memory decay & memory use probability based on...<br>1) distance between patches<br>2) time since last visit to the patches |
| Thompson et al. 2022 | Brown Bear | Current resource quality + time since the last visit | Forage | Hidden Markov Model with SSF Framework<br>Likelihood of animals moving in non-stationary states with a cognitive map | Maximum-likelihood estimates (MLE) | 1) distance from turbid water to riparian areas where bears selected<br>2) vegetation classes<br>3) density of ground squirrels and alpine sweet vetch<br>4) distance from human settlements |
| Oliveira-Santos et al. 2016 | Feral hog | Utilization distribution of an animal's trajectory | Home range | SSF framework with an OD as a response variable in the model | Step Selection Functions (SSFs) with conditional logistic regression | 1) Land cover types<br>2) Time of day<br>3) BRB densities<br>4) Residence time |
| Northrup et al. 2016 | Mule deer | Differences between animals' occurrences throughout time | Home range | SSF framework with an OD as a response variable in the model | Hierarchical Bayesian regression (MCMC) | 1) Tree<br>2) density of drilling well pads<br>3) average NDVI difference<br>4) major road density<br>5) terrain ruggedness<br>6) fat<br>7) age |
|  |  |  |  |  |  | 8) natural gas facilities' density<br>9) total snow depth difference<br>10) pipeline density difference |
| Schlägel et al. 2017 | Wolf | Time since the last visit | Territorial defense | Spatially explicit random walk model similar framework of...<br>+ selection-free movement kernel + selection functions | Maximum-likelihood estimates (MLE) | 1) Prey density<br>2) Distance to territory boundary |
| Ranc et al. 2021 | Roe deer | Food availability | Home range | Likelihood of animals moving with cognitive selection functions | Maximum-likelihood estimates (MLE) | 1) Min. daily temperature<br>2) within state resource access<br>3) between-state resource access<br>4) illumination index (dawn & dusk)<br>5) changes in the illumination index |
| Ranc et al. 2022 | Roe deer | Surrounding available habitats | Home range | Likelihood of animals moving with cognitive selection functions | Maximum-likelihood estimates (MLE) | 1) Landcover - reforested<br>2) Landcover - agriculture<br>3) step lengths<br>4) Memory<br>5) Memory decay |
| Gurarie et al. 2022 | Wolf | Predation success & Territorial marking | Territorial movement | A discrete choice modeling framework | Maximum-likelihood estimates (MLE) | 1) selected zone per each trip<br>2) predation quality<br>3) boundary coverage<br>4) mass of prey item<br>5) time since predation events<br>6) number of kills<br>7) boundary sizes |
| Merkle et al. 2019 | Mule deer | Differences between animals' occurrences throughout time | Migration | SSF framework with a distance and turning angle bias as parameters in the model | Step Selection Functions (SSFs) with conditional logistic regression | 1) Elevation<br>2) % tree cover<br>3) Distance to roads<br>4) Terrain position index<br>5) Integrated NDVI<br>6) Rate of Green-up<br>7) Distance to the previous route<br>8) Direction to the previous range |

Popular analytical frameworks, such as integrated Step-Selection Analyses (SSAs) (Avgar et al., 2016; Fieberg et al., 2021; Fortin et al., 2005; Thurfjell et al., 2014) have been used to identify signals of memory from such observations (Merkle et al., 2019; Oliveira-Santos et al., 2016; Rheault et al., 2021; Ran et al., 2022; Schlägel & Lewis, 2014). We focus our review on SSAs because of their flexibility and ease of use due to readily available statistical software (Signer et al. 2019), but also because of their continuous methodological development (Klapstein et al. 2024, Michelot et al., 2024; Potts & Börger., 2023; Signer et al., 2024). Nonetheless, several challenges remain before these approaches can be widely adopted. These include technological challenges associated with managing tracking data, creating models with different “familiarity” or “memory” covariates, and drawing biologically accurate conclusions from these models. Here, we provide an overview of methods for parameterizing memory effects in SSAs to help guide practitioners wishing to identify or quantify the effects of memory on animal movement. We also offer several examples with annotated code and then discuss the strengths and limitations of current approaches and future directions for memory-informed movement research.

## 2. Exploring how memory influences animal movements using SSAs

SSAs are widely used to quantify influences on animal movement (Avgar et al., 2016; Fieberg et al., 2021; Fortin et al., 2005; Thurfjell et al., 2014). SSAs model movement in discrete time using two model components: 1) a selection-free movement kernel describing how animals move in the absence of habitat selection, defined using distributions of step length and turning angle, and 2) a selection function describing animal preferences concerning the habitat attributes at each step’s endpoint (Box 1). Model parameters in SSAs can be estimated using commonly available statistical software that implements conditional logistic regression (Avgar et al., 2016; Fieberg et al., 2021). Because they allow one to model and predict dynamic space-use patterns using accessible and available software, SSAs are attractive to movement ecologists and are widely used to analyze animal tracking data (Signer et al., 2019). Further, SSAs have strong connections to other popular methods for modeling animal movement; they have been shown to be equivalent to biased correlated random walks (Duchesne et al., 2015) and can be approximated by diffusion-taxis models (Potts & Schlagel. 2020). In addition, certain continuous time movement models can be recast as SSAs (Eisaguirre et al., 2024).

A variety of spatiotemporal familiarity covariates can be included in an SSA to model the effects of memory (Box 2; Box 3). For example, one may estimate an occurrence distribution (OD; Fleming et al. 2015, Alston et al. 2024) describing the relative use of the landscape over a specific period in the past (e.g., a continuous surface of either the relative intensity of use or a binary presence/absence variable). By including the OD as a spatial predictor in an SSA, one can evaluate whether an animal’s current landscape usage is biased toward or away from previously visited locations (Rheault et al., 2021; Oliveira-Santos et al., 2016; Wolf et al., 2009). A notable limitation of this approach is the risk of overestimating the importance of familiarity due to unaccounted-for habitat attributes (e.g. if the OD reflects the distribution of an unobserved resource; see Picardi et al. 2023). The OD approach further necessitates that users choose an appropriate time window in the past for calculating the OD (Figure 1). Using a limited time window for calculating the familiarity covariate implies an abrupt memory decay function where past locations are memorized for a fixed amount of time and then forgotten completely.

**Figure 1.**
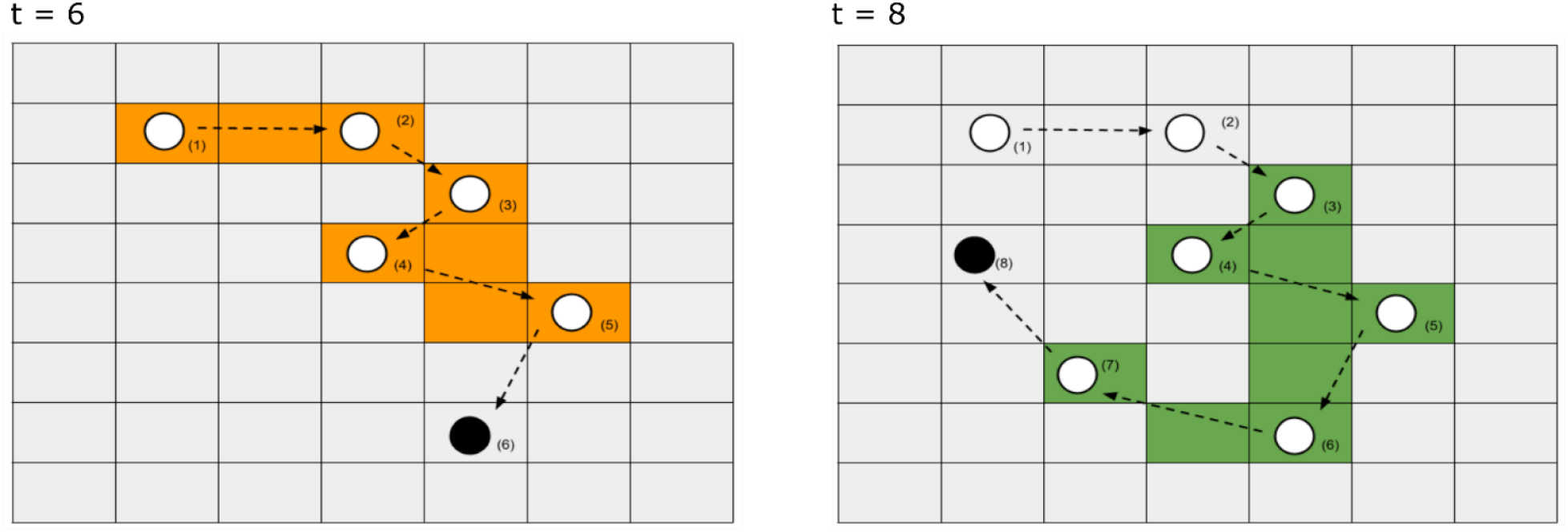
Spatial memory can be quantified using an animal’s occurrence distribution (OD) measured over some prior period (colored areas: 5-day periods at time 6 and time 8 = orange vs green) which captures an animal’s movement path and its uncertainty. A time-varying covariate can be constructed by updating the OD at regular time intervals. This updating step ensures that distant experiences are eventually forgotten and no longer play a role in driving animal movement. Users must choose an appropriate time window in the past for calculating the memory covariate and how often to update it.

Alternatively, one could choose to continually update the OD from the first to the last location, which would imply the animal never forgets its past experiences. With either approach, the OD effectively weights all previously visited areas within the specified time window equally, regardless of how long ago the animal visited the location. Another option is to allow more recent (or distant) memories to have more influence on current movements by replacing the OD with a covariate representing the length of time since the animal last visited a location (TSLV). Or, one can create multiple ODs reflecting space use during the recent or the more distant past and allow the model to determine optimal weights given to short-term and long-term memories represented by these covariates (e.g., Oliveira-Santos et al., 2016).

For migratory species that navigate relatively long distances, familiarity predictors could include distances between current and previous migratory trajectories or angular covariates that compare the direction of an animal’s movement in relation to a previous seasonal range (Figure 2). These methods can describe an animal’s use of memory for navigation and capture its tendency to use familiar migration routes and consistent but seasonally varying home ranges (Merkle et al., 2019).

**Figure 2.**
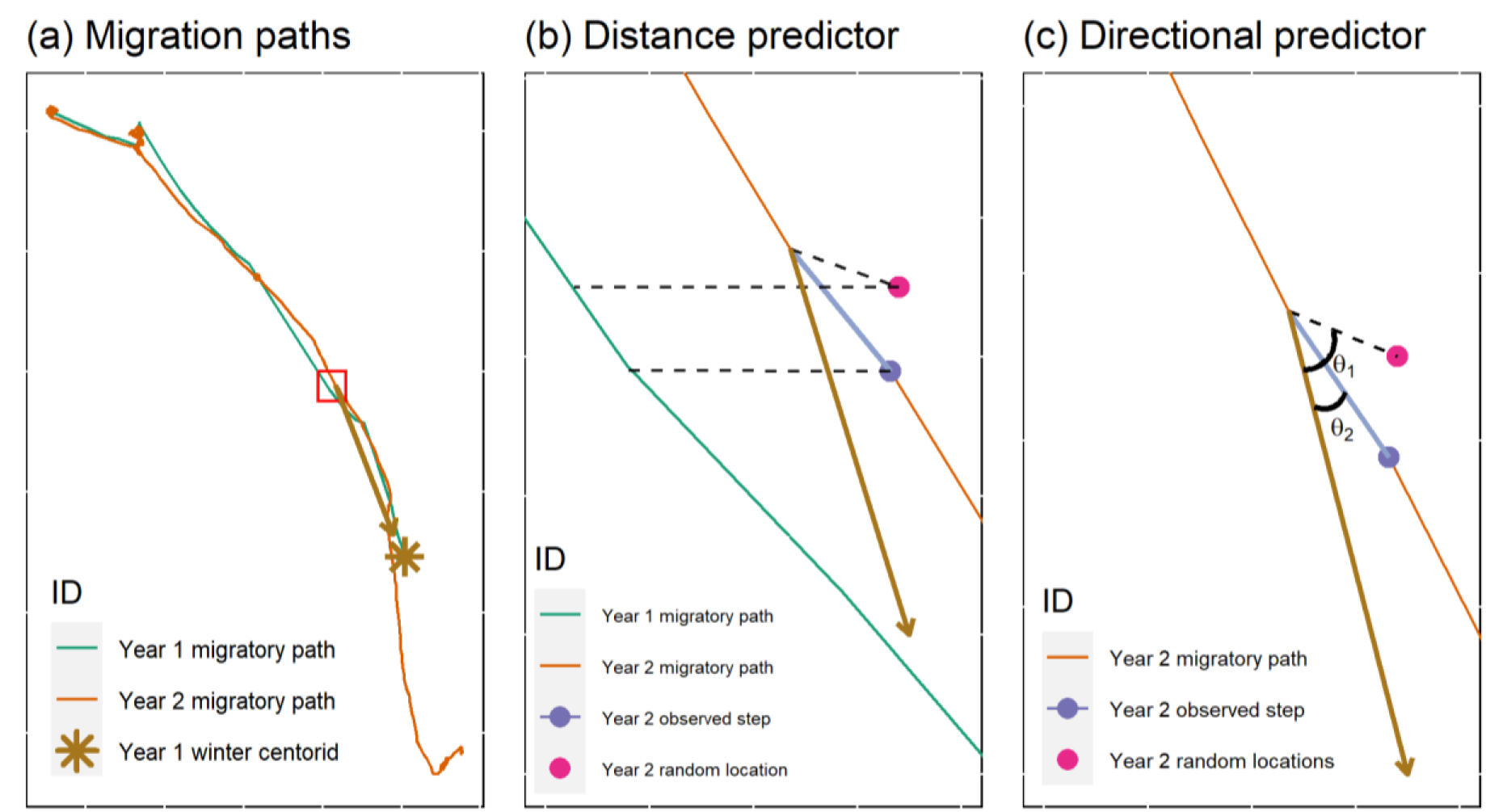
Familiarity covariates used in applications involving migratory animals. The red bounding box in panel (a) displays the area zoomed into for boxes b and c. Memory can be quantified using a distance predictor calculated as the minimum distance between current and past migratory paths (panel b). Specifically, we can calculate the distances (dashed lines) between Year 2 (observed [navy], random [pink]) locations and Year 1 (green) migratory paths. Memory can also be included as a directional bias predictor by comparing whether the current year’s steps are biased toward the previous season’s range (panel c). This bias predictor can be calculated using the angles, *θ*_1_,*θ*_2_, between the step (observed [navy], random [pink]) and the centroid of the previous year’s winter range.

Unless it is reasonable to assume that the animal lacked any memory at the onset of tracking, it is necessary to ‘sacrifice’ some early positional data to calculate the familiarity predictor. For some animals, including those that have long lifespans or live in highly seasonal environments, the ‘memory build-up’ period may need to be one year or more, unless there are reasons to believe memory (or its effect) decays over much shorter time scales. For example, Avgar et al. (2015) used a full year of ‘memory build-up’ data to model woodland caribou (*Rangifer tarandus caribou*) space-use patterns over a subsequent year but have found no indication of memory decay within that time (indicating the need to use an even longer ‘memory build up’ period). Alternatively, studies of young or dispersing individuals and data from translocation experiments offer opportunities to model memory formation as individuals enter new and unfamiliar environments and learn how to navigate them efficiently (Dalziel et al., 2008; Wolf et al., 2009; Fagan et al., 2013; Lewis et al., 2008; Ranc et al., 2021. 2022; Aikens et al., 2024; Brønnvik et al., 2024). However, datasets containing movements of individuals in unfamiliar landscapes are rare and difficult to obtain due to the high cost of translocation experiments and a tendency to avoid tracking juveniles due to often high mortality rates (Sergio et al., 2011; Weise et al., 2014).

## 3. Case studies

In this section, we review memory-informed movement models for animal tracking data using 4 case studies. The first two examples, involving data from sandhill cranes (*Antigone canadensis*) and feral hogs (*Sus scrofa*), demonstrate how one can use a spatial familiarity predictor calculated from a past OD to explore whether animals retain information from their past experiences and for how long. The third example considers the migratory movements of mule deer and illustrates the use of two familiarity predictors formed using the minimum distance between the current and last year’s migratory paths and the cosine of the angle between the direction of a current movement step and the previous year’s centroid of locations. The last example, involving data from a brown bear, uses a spatiotemporal covariate to quantify how long it has been since the individual last visited spatial locations, and thus, how memory may influence revisitation rates. We provide a workflow to reproduce the main components of the habitat- and memory-based SSF from all the case studies, demonstrating the relative importance of memory-based and habitat metrics in an SSF of animal movement (see Appendix codes for the case studies). We also highlight challenges and decisions that the user must make when applying these methods to their tracking data (Box 2).

## 1. Sandhill Crane – ‘fixed-time’ OD

Sandhill cranes breed throughout North America during the summer and migrate south for the winter. During their first year, juveniles migrate to overwintering areas with their parents and then disperse from the family group either during the spring migration or upon arrival to the natal territory the following spring (Hayes, 2015; Tacha et al., 1989). During the first few years of independence, subadult cranes typically make long-distance movements across the landscape during the summer; in contrast, the movements of breeding adults are largely constrained to their breeding territories (Wolfson et al., 2020). Once cranes become successful breeding adults, typically between 4-6 years old, they use their accumulated knowledge of the landscape to return and nest in the same breeding area in subsequent years (Nesbitt, 1992; Tacha et al., 1989).

As an illustrative example of how a sandhill crane’s space use constricts each year as the crane learns its landscape and develops a breeding territory, we consider a 5-year dataset of global positioning system (GPS) telemetry locations of a sandhill crane (with a 15-minute fix interval), starting from the time of fledging (Wolfson, 2018; Wolfson et al., 2020). A visualization of summer locations shows that the spatial coverage visited by the crane decreases each year as it ages, selecting locations it had previously visited (Figure 3a). This pattern suggests that the crane may be using its past experience to decide where to establish a breeding territory.

**Figure 3.**
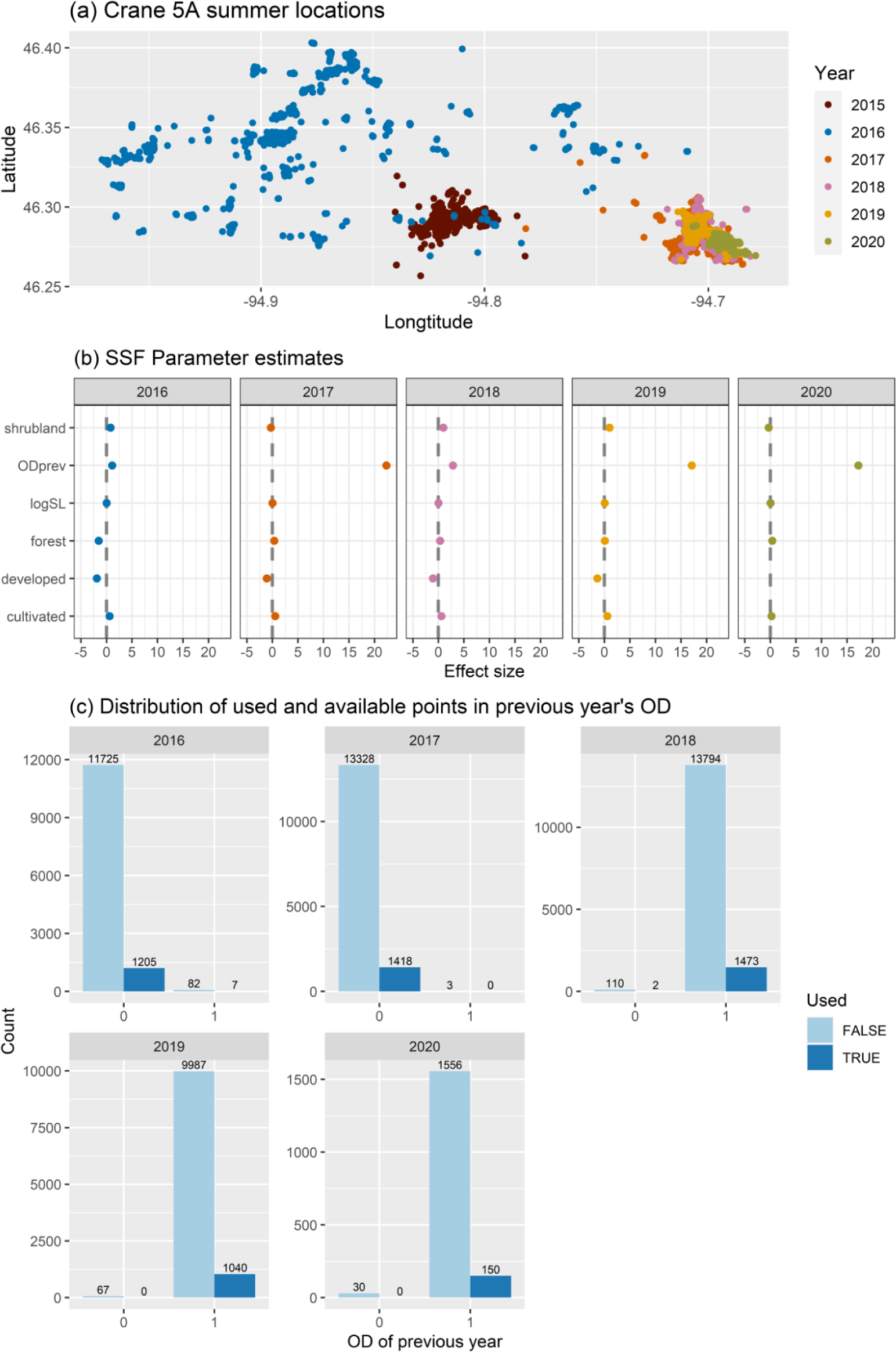
Visualization of Sandhill crane tracking data and coefficients from fitted step-selection functions with a memory covariate formed using an OD capturing previously visited locations **a)** Movement patterns of an individual sandhill crane during summer seasons (06/15-09/22) from 2015 to 2020. **b)** Parameter estimates including those for the memory covariate (Odprev: OD from the previous year). **c)** Distribution of each year’s used and available locations for sites (i.e., grid cells) that were (OD = 1) and were not (OD = 0) visited in the previous year. Numbers at the top of the bars indicate the number of locations in each group.

To quantify the potential effects of familiarity on the crane’s summer movements, we calculated areas associated with the 95% contour of an OD estimated from the previous year’s summer locations using the probability density functions from a continuous-time movement model conditioned off the Brownian bridge (Fleming et al., 2015). We then fit a separate SSF to each year of data, including the previous year’s OD as a predictor in the model. A positive coefficient associated with the previous year’s OD would suggest the crane prefers to revisit sites it visited in the previous year (compared to *equally accessible and otherwise identical sites* that it did not visit during the previous year, where accessibility is determined by the selection-free movement kernel; Fieberg et al. 2021). If the coefficient is negative, the opposite interpretation holds.

Coefficients associated with the previous year’s OD were positive in all years (Figure 3b), even though the crane rarely revisited previously used locations in the first two years (Figure 3c). Interpretation of the coefficients associated with categorical predictors can be difficult, as they reflect a ratio of ratios (use: availability ratio for one class versus the use: availability ratio of a reference class; Fieberg et al. 2021). The positive coefficients for the previous year’s OD in 2016 and 2017 reflect the fact that the use-availability ratio associated with grid cells visited in the previous year was higher than the use: availability ratio for grid cells that were not visited in the previous year. Thus, we end up with a positive coefficient for the OD predictor even though most of the areas encountered and used by the crane represent areas that were not visited in the prior year. Recall our interpretation of a positive coefficient, *if presented with two equally accessible locations* (same step length and turn angle required to reach both locations) that only differ in whether they had been visited in the previous year (i.e., they had the same landcover class), the crane would be more likely to select the location it was familiar with. A limitation of this model and application is that the crane was rarely presented with this type of choice. In the first two years, it was rarely found near sites it had previously visited whereas in years 3-5 it rarely visited areas it was unfamiliar with. This example highlights a limitation of the OD approach and the need for new methods that can capture tradeoffs in exploratory and informed movements as young individuals learn to navigate the landscape. Another limitation is that the birds may be selecting for an unobserved resource, one that we did not include in our analysis, which introduces a hidden correlation between their past and current space use.

## 2. Feral Hog – OD with temporal variation: short- and long-term memories stratified by time of day

Hogs were introduced to the Pantanal wetland about 300 years ago, and currently represent the highest wild mammal biomass in this region. They are crepuscular-nocturnal, social, long-lived, cooperative animals that forage at the edges of water bodies and in ephemeral pools that become increasingly rare during the winter dry season. Hogs lack sweat glands and behaviorally mitigate heat stress by spending the hot hours of the day resting in forest patches.

As animals move, they can access and update their reference memory (long-term acquisition) and working memory (short-term acquisition) to navigate through space. Still, the stored spatial information may or may not be used depending on the current animal needs and context, which generates temporal heterogeneity (e.g., within the day, and seasons) in the use of that information. Oliveira-Santos et al. (2016) modeled the hidden process underlying spatial memory acquisition using spatiotemporal covariates generated from Biased Random Bridge kernel density estimates based on residence time (Benhamou, 2011). Specifically, for each individual step, they used previous locations to build multiple spatial maps, formed using different time windows, that could affect the hogs’ future movement decisions. These maps were then continuously updated as the individual moved through the landscape.

Oliveira-Santos et al. (2016) considered 4 conceptually different hypotheses regarding how hogs process and use spatial information, combining long- and short-term memory with differential use of stored memory within the daily cycle. To represent long-term memory, familiarity covariates were constructed and constantly updated with all previous locations as the individual moved, whereas short-term memory was represented using a spatiotemporal covariate that kept track of just the last 3 days. Additionally, long- and short-term familiarity covariates were also built considering only daytime or only nighttime locations, which they referred to as long-term temporal and recent-temporal memory, respectively. When fitting models for these last two cases, movement steps taken at night or during the day were paired with familiarity covariates generated from previous locations collected only at night or day, respectively.

All tracked hogs strongly selected for previously visited areas, mainly those associated with short-term memory. Most of these individuals (65% of the tracked hogs) appeared to use working memory as part of their movement process, as covariates generated from recent nocturnal locations were better at predicting future nocturnal use than covariates generated from all time periods. Importantly, the effect of familiarity also varied within the day, being more important during the daylight hours when individuals were sleeping in well-known places than at night when animals were foraging and were more willing to take risks by walking through less familiar areas. Although hogs are acknowledged to have high cognitive skills and memory retention, Oliveira-Santos et al. concluded that they relied mainly on recent spatial information because the distribution of prime food resources in the study area responds quickly to foraging pressure and changes in water levels.

## 3. Mule Deer – Migratory paths and angles

Many large ungulates are migratory, capitalizing on seasonal and spatial variation in food, predation, and hospitable conditions (Avgar et al., 2013). Mule deer are a concentrate forager (i.e., prefer to consume high-quality food) that display some of the longest terrestrial migrations in North America (Joly et al., 2019), spending winters in arid, low-elevation sagebrush, grassland, and desert ecosystems, and then migrating (up to 400 km) into montane ecosystems at higher elevations for summer.

Unlike some other migratory species (e.g., Sierra Nevada Bighorn Sheep; Berger et al., 2022), Mule deer migrate well outside their perceptual range (e.g., what they see, hear, and smell at a given moment), yet they display relatively strong fidelity to seasonal ranges (Morrison et al., 2021) and migration routes (Sawyer et al., 2019). On average, mule deer migrate on the same path during spring year after year 81% of the time (Sawyer et al., 2019). In some cases, such migrations occur across relatively vast expanses of flat deserts, rolling hills, and thick forests, where sensory abilities such as vision may provide limited cues for navigation (Sawyer et al., 2016).

Evidence suggests that mule deer may navigate during migration by memorizing the path of their previous migration route and the general location of their seasonal ranges. The relative influence of these memorized spatial locations can be assessed in a movement model that first considers several other habitat features that may influence ungulate movement and space use.

Merkle et al. (2019) examined the relative role of memory usage versus local variation in habitat on mule deer navigation during migration. They found that variables indexing past experience (distance to the previous migration route and direction to the previous seasonal range) were 2 to 28 times more predictive of migratory movements than local variation in habitat. Alternative explanations for these long-distance migrations include following scent trails or other conspecifics, however, those explanations are not well supported due to the fact that these deer spend on average 81% of their migration walking on the exact same path as in the previous year (Sawyer et al. 2019).

## 4. Brown Bear – time since last visit (TSLV)

Brown bears are opportunistic omnivores found in North America, Europe, and Asia (Pasitschniak-Arts, 1993). Their life history strategies and dietary compositions vary greatly depending on the environment they live in (Finnegan et al., 2021; García-Rodríguez et al., 2021; MacHutchon & Wellwood, 2003), but their ability to navigate towards previously visited food patches is ubiquitous throughout their natural range (Selva et al., 2017; Thompson, Lewis, et al., 2022; Wirsing et al., 2018). The “barren-ground grizzly bears” found in the Canadian Arctic are unique for many reasons, including their foraging and denning behavior (Edwards & Derocher, 2015; McLoughlin, 2002). These bears’ food resources are only available for a short, albeit predictable, portion of their active seasons (Edwards & Derocher, 2015), so in addition to returning to the correct spatial location where food was present, bears must identify the temporal pattern of this resource and revisit the patch at the correct time. Ecologists interested in understanding how these bears incorporate memory into their movement patterns must implement models that account for these nonlinear temporal dynamics.

We modified the model developed by Schlägel & Lewis (2014) so that it could be implemented in the step-selection framework and fit using conditional logistic regression. This required using distributions from the exponential family (gamma, von Mises) to model the distribution of step lengths and turn angles, respectively (Avgar et al. 2016; Fieberg et al. 2021). In addition, we included both linear and quadratic terms to model the influence of time since last visit (TSLV) on the movement patterns of an adult female brown bear in the Mackenzie River Delta region of the Northwest Territories, Canada. This bear was immobilized and fitted with a GPS collar that recorded its location every 4 hours during the active season (the time in which the bear was out of its den).

We calculated TSLV as the difference between the current timestamp and the last time the bear visited a series of 2 x 2 km grid cells, updating the map at each observation time. This familiarity covariate allowed “revisitation” to occur when the animal was in the perceptual vicinity of an area it previously visited (i.e., within the same grid cell), without necessitating that the animal returns exactly to its previous coordinates (Schlägel & Lewis, 2014). We discarded the first year of location data as a “burn-in”, and set TSLV to 365 for any grid cells where TSLV was missing, effectively assuming that these cells had been visited just prior to the first observed location. The need for a burn-in period is a limitation of this approach but is necessary since we have no history of the bear’s past visits prior to the start of data collection. A sensitivity analysis can be performed to evaluate whether results change if the length of the burn-in period is increased or decreased. To analyze how selection strength varied with TSLV, we calculated the relative selection strength (RSS) at different TSLV values (Avgar, 2017). As with other habitat selection models, by including both linear and quadratic terms, we were able to identify a non-linear response to TSLV with intermediate TSLV values of approximately 350 days displaying the strongest selection (Figure 4). These results agree with those of Thompson et al. (2022b), who also found that bears tended to revisit sites seasonally.

**Figure 4.**
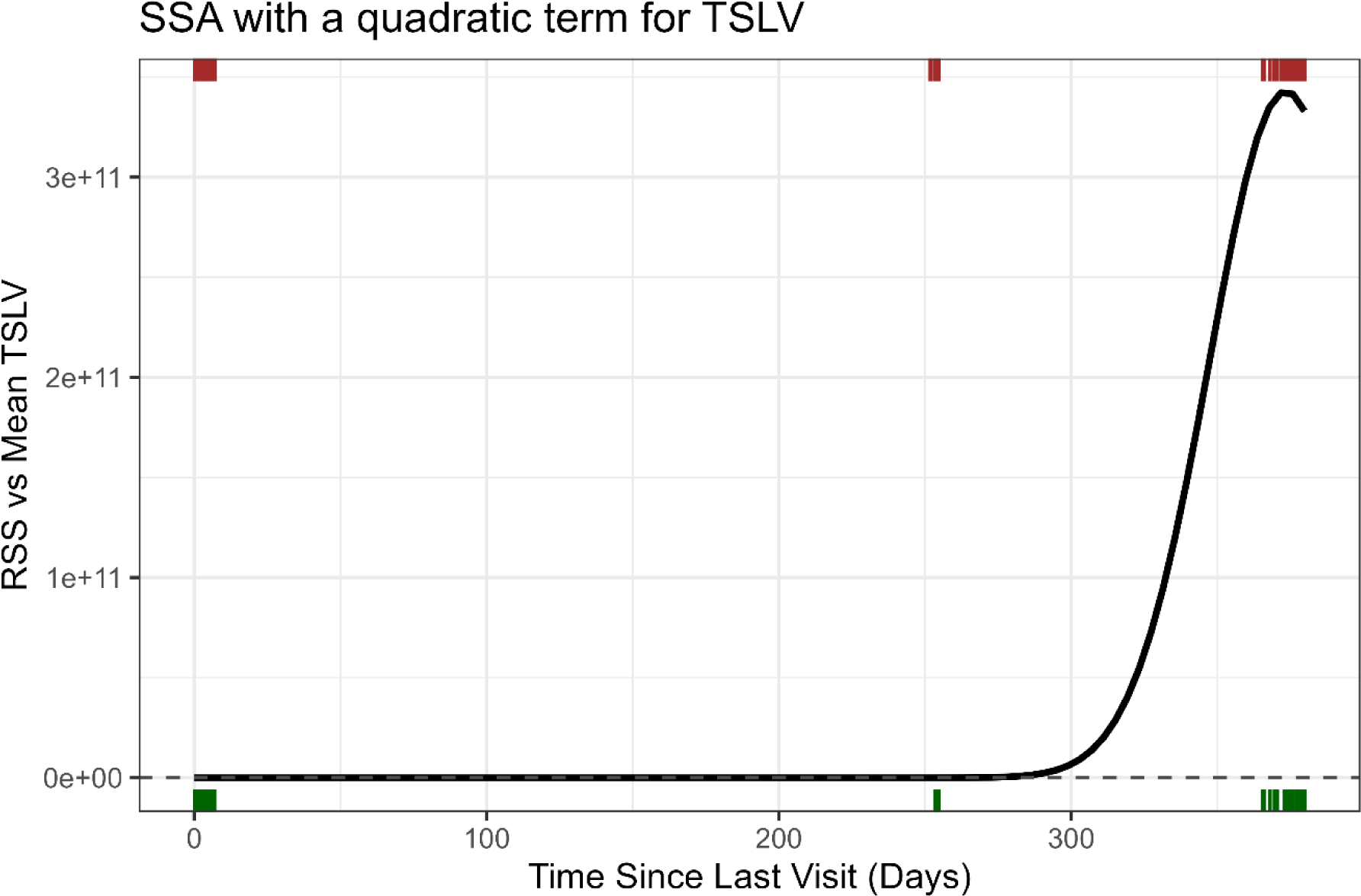
Visualization of the relative attractiveness of previously visited locations as a function of time since last visit (TSLV) modeled using linear and quadratic terms in a step-selection analysis of brown bear location data with the distribution of used (green, bottom) and available (brown, top) locations shown along the x-axis.

Although the TSLV approach can capture the potential memory effects of wildlife over time, the interpretation of the model may depend on the user’s choice of resource covariates. For example, we used berries as a seasonal resource covariate. However, the importance of the memory covariate (TSLV) might change if additional environmental covariates are added to the model (Picardi et al., 2023). Similar to Merkle et al. (2019), Thompson et al. (2022b) compared their models with memory to “resource-only” models without memory, finding that the former models better fit the data. Yet, it is nearly impossible to perfectly quantify the distribution of food resources on the landscape, and “resource-only” models might have performed better if more environmental data could be acquired. Even when alternative models are considered and compared rigorously to a “memory model”, it is important to consider how these alternative models may fall short in quantifying their desired hypotheses.

## 4. Discussion

Modern GPS tracking systems generate massive telemetry datasets by following individual animals over a long time on a global scale. With this abundance of available data, it is now possible to develop models that evaluate how memory relates to animal movement (Nathan et al., 2022), which has inspired the recent development and application of many such models (see Table 1). Our review focused on approaches that account for familiarity with different areas of the landscape by including spatial, spatiotemporal, or angular covariates as predictors in step-selection analyses. These frameworks can be implemented using available statistical software for fitting conditional logistic regression models, which we demonstrate using multiple tracking data sets and annotated code examples. Although some data development is necessary before fitting the models (e.g., to create the memory predictors and generate random steps), users can leverage R packages to make these steps easier. For example, the ctmm package (Calabrese et al., 2016) can be used to calculate ODs, and the amt package (Signer et al., 2019) can be used to generate random steps necessary for parameterizing the model.

Individual-based models of animal movement are increasingly used to inform conservation and management at the population or even species levels (Ariello et al., 2023; Hofmann et al., 2023; Whittington et al., 2022). Prime among these applications is the use of habitat-selection models to identify critical habitats, delineate range boundaries, and project spatial distribution across space or time (Osipova et al., 2019). A strong selection for certain habitats or environmental features and conditions may occur with or without memory, and it is important to consider how models that incorporate familiarity covariates may alter inferences. On the one hand, we might expect mild to moderate collinearity between familiarity covariates and other important environmental drivers, which can make it challenging to quantify their unique contributions (Picardi et al., 2023). On the other hand, accounting for familiarity and memory effects should reduce bias associated with estimators of habitat-selection strength.

Consider an animal that is both attracted to certain habitats and to places it has used in the past; if we ignore the latter, we will consequently overestimate the former, putting more weight on habitat attraction than it truly has (Picardi et al., 2023). Similarly, it is important to consider the effect of environmental drivers when looking for influences of memory on animal movements (Van Moorter et al., 2013). Memory-free movement models that allow for habitat selection have also been suggested as null models for evaluating evidence of memory (Picardi et al., 2023). An additional benefit to considering memory effects in SSAs is the option to simulate these effects under various management scenarios (Forrest et al., 2024). For example, managers might be interested in assessing the likelihood of successfully relocating an animal and having it establish a home range in a new site (Ranc et al., 2021; Berger et al., 2022). Explicitly modeling the process of building and responding to increasing knowledge of the landscape may be critical for obtaining reliable predictions.

Information gathering, an essential prerequisite for memory, is governed by sensory ecology (Dusenbery., 1992, Dukas., 1998; Dall et al., 2005; Jordan & Ryan., 2020; Dominoni et al., 2020;), and it is in this context that we must consider the limitations associated with the use of SSA to infer memory. The information available to the animal about any given spatial location (the ‘signal’) is a function of the animal’s position in relation to that location (the source of the signal), the strength of the signal (e.g., the intensity of odor, light, or sound), the overlap with similar signals coming from elsewhere on the landscape, and the animal’s sensory capacity to perceive and process the signal (which in itself may be a complex function of the animal’s morphology and physiology). Information can only be committed into memory if it is being perceived. Moreover, it is very likely that the weight given to memorized information (or its retention time) is modulated by the signal-to-noise ratio at which the information was perceived. The models we described here assume, for the most part, that animals sense and retain information from only ‘visited localities’, which are arbitrarily sized spatial units (typically corresponding to a single pixel of the available environmental data). Furthermore, most SSFs only assess what the animal can perceive within local ranges (i.e., what is nearby), though angular and distance-to-covariates allow modeling perception over larger spatial scales. Thus, SSFs are a useful, but extreme simplification of the true underlying sensory ecology.

More mechanistically inclined frameworks for explicitly modeling the sensory processes involved in memory buildup have been proposed (e.g., Avgar et al. 2013, 2015; Merkle et al., 2014; Ranc et al. 2022; Thompson, Lewis, et al. 2022); however, fitting these models to wildlife tracking data is still a challenge. These frameworks typically include additional free parameters used to construct spatiotemporal covariates that determine how perceived habitat quality varies over time (e.g., by modeling how perception decays with distance from the individual, and memory decays temporally). As a result, it is not possible to fit these models using standard statistical software developed for SSAs. Instead, the parameters governing the spatiotemporal covariate must be estimated simultaneously with other habitat-selection and movement parameters through a custom-written likelihood function that can be optimized using Markov chain Monte Carlo or other numerical optimization methods. We include an example from Thompson et al. (2021) in our supplementary material to demonstrate this approach. Methods that more realistically model the process of memory formation should be pursued, but we suspect that most practitioners will continue to explore the role of memory on animal movements using simpler models that can fit within the standard SSA framework using conditional logistic regression. The strength of this approach is that it can be easily and widely applied to tracking data. Still, these efforts should be complemented by additional experiments and more realistic mechanistic models to better understand the multifaceted ways that animals use memory to navigate their landscapes (Ranc et al., 2022; Wild et al., 2023).

## Contributions

DK and JF conceived of the topic, and DK led the writing of the manuscript. Case studies were led by DK, DW, and JF (sandhill crane), DK, PT, and JF (brown bear), JM (mule deer), and L O-S and JDF (hog). All authors discussed, edited, read, and approved the final manuscript.

## Ethics declarations

### Ethics approval and consent to participate

Not applicable.

### Consent for publication

Not applicable.

### Availability of data and materials

The datasets and code associated with the case studies would be archived in the Data Repository for the University of Minnesota (DRUM) if the manuscript is accepted.

### Box 1. Step-Selection Analyses (SSAs)

Step-Selection Analyses (SSAs) model the conditional probability, *p*(*s_t_ | H_t_*_-1_;*β_m_β_w_*), of finding an individual at a location *s_t_* at the time *t* given a set of previously visited locations,*H_t_*_-1_, using a selection-free movement kernel, *k*(*s_t_ | H_t_*_-1_;*β_m_*), which describes how animals would move in the absence of habitat selection, and a movement-free habitat-selection function, *w*(*s_t_; t,β_w_*), which describes the animals’ preferences for certain environmental features (e.g., variables representing resources, risks, and or other conditions; Matthiopoulos et al. 2023):

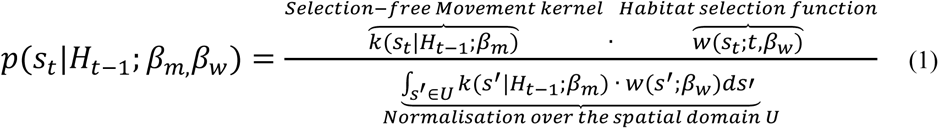

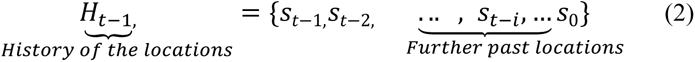

*β_m_* contains parameters in the step-length and turn angle distributions (*β_m_*_1_,…,*β_mq_*), and *β_w_* contains resource- selection parameters that quantify the attractiveness of different locations using a vector of selection coefficients (*β_w_*_1_,…,*β_wp_*) for each environmental covariate (*r*_1_(*s_t_*),…,*r_p_*(*s_t_*)). *s′* ∈ U describes all the locations within the spatial domain *U*. To calculate a step length, *sl*, two locations are required, (*s_t_*, *s_t_*_-1_). Similarly, a turning angle, *ta*, is calculated using the current and the past two locations (*s_t_*, *s_t_*_-1_, *s_t_*_-2_).

### Box 3. Familiarity function in SSFs

Movement ecologists have described familiarity with different parts of the landscape that animals experienced using familiarity covariates, which we formalize via a familiarity function, *f.* Including an exponential familiarity function allows the attractiveness of different locations to be governed by both environmental and familiarity covariates within the traditional SSF framework 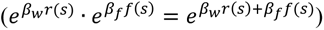. Examples of familiarity covariates include occurrence distributions (ODs) reflecting the intensity of past space use in the study area, time since last visit (TSLV), migratory distances between current and previously used paths, and angular covariates used to capture bias toward previously used migratory ranges:

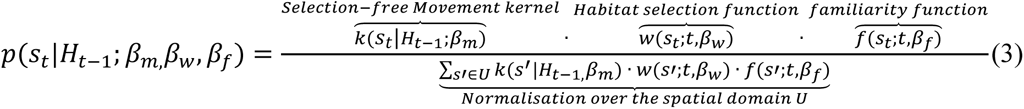

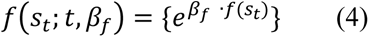

**Box 2.** Guide for the memory-informed movement modeling within SSF framework.

| Question | Biological example | Method | Caveats, challenges, limitations | Implementation parameters or choices |
| --- | --- | --- | --- | --- |
| Are animals more likely to visit areas that are more familiar to them? | Feral hogs primarily rely on recent experiences when making night time foraging decisions (Oliveira-Santos et al., 2016). | Occurrence Distribution (OD) | The effects of unmodeled environmental drivers may be wrongly attributed to memory and vice versa (Picardi et al. 2023). | Time window for measuring past space use.<br>Length of burn-in periods (how much early positional data to sacrifice to estimate the familiarity predictor?) |
| Do animals prefer to visit areas they have not visited recently? (and if so, is there an optimal return time based on foraging or predation experience?) | Predators like wolves and brown bears delay returning to previously visited kill sites so that prey numbers may rebound to normal levels (Gurarie et al., 2022; Selva et al., 2017). | Time Since Last Visit (TSLV) | The effects of unmodeled environmental drivers may be wrongly attributed to memory and vice versa (Picardi et al. 2023).<br>TSLV may be relatively homogeneous within an animal's movement kernel. | "Patch" size (what is the typical spatial grain of the animal's cognitive map?)<br>Length of burn-in periods (how much early positional data to sacrifice to estimate the familiarity predictor?) |
| Do migratory animals use consistent migration routes to return to the same seasonal ranges each year? | Mule deer (*Odocoileus hemionus*) have been shown to use the exact same migration routes and seasonal ranges, likely relying on their memory of those spatial locations to help them navigate (Merkle et al., 2019). | Distance and directional bias toward previously used areas across large spatial scales | The effects of unmodeled environmental drivers and social cues may be wrongly attributed to memory and vice versa (Picardi et al. 2023). | Whether to use angular covariates or distance-to-covariates.<br>Time window for measuring past space use. |

## References

Aarts, G., Mul, E., Fieberg, J., Brasseur, S., van Gils, J. A., Matthiopoulos, J., & Riotte-Lambert, L. (2021). Individual-level memory is sufficient to create spatial segregation among neighboring colonies of central place foragers. The American Naturalist, 198(2), E37–E52. 10.1086/715014

Abrahms, B., Hazen, E. L., Aikens, E. O., Savoca, M. S., Goldbogen, J. A., Bograd, S. J., Jacox, M. G., Irvine, L. M., Palacios, D. M., & Mate, B. R. (2019). Memory and resource tracking drive blue whale migrations. Proceedings of the National Academy of Sciences, 116(12), 5582–5587. 10.1073/pnas.1819031116

Aikens, E. O., Nourani, E., Fiedler, W., Wikelski, M., & Flack, A. (2024). Learning shapes the development of migratory behavior. Proceedings of the National Academy of Sciences, 121(12), e2306389121.

Aiello, C. M., Galloway, N. L., Prentice, P. R., Darby, N. W., Hughson, D., & Epps, C. W. (2023). Movement models and simulation reveal highway impacts and mitigation opportunities for a metapopulation-distributed species. Landscape Ecology, 38 (4), 1085–1103. https :465//doi.org/10.1007/s10980-023-01600-6

Alston, J. M., Fleming, C. H., Noonan, M. J., Tucker, M. A., Silva, I., Folta, C., … & Calabrese, J. M. (2022). Clarifying space use concepts in ecology: range vs. occurrence distributions. BioRxiv, 2022-09.

Avgar, T. (2017). Relative selection strength: Quantifying effect size in habitat-and step-selection inference. Ecology and Evolution, 7(14), 5322–5330.

Avgar, T., Baker, J. A., Brown, G. S., Hagens, J. S., Kittle, A. M., Mallon, E. E., McGreer, M. T., Mosser, A., Newmaster, S. G., Patterson, B. R., Reid, D. E. B., Rodgers, A. R., Shuter, J., Street, G. M., Thompson, I., Turetsky, M. J., Wiebe, P. A., & Fryxell, J. M. (2015). Space-use behavior of woodland caribou based on a cognitive movement model. Journal of Animal Ecology, 84(4), 1059–1070. 10.1111/1365-2656.12357

Avgar, T., Deardon, R., & Fryxell, J. M. (2013). An empirically parameterized individual based model of animal movement, perception, and memory. Ecological Modelling, 251, 158–172. 10.1016/j.ecolmodel.2012.12.002

Avgar, T., Potts, J. R., Lewis, M. A., & Boyce, M. S. (2016). Integrated step selection analysis: Bridging the gap between resource selection and animal movement. Methods in Ecology and Evolution, 7(5), 619– 630. 10.1111/2041-210X.12528

Bauer, S., & Hoye, B. J. (2014). Migratory Animals Couple Biodiversity and Ecosystem Functioning Worldwide. Science, 344(6179), 1242552. 10.1126/science.1242552

Benhamou, S. (2011). Dynamic Approach to Space and Habitat Use Based on Biased Random Bridges. PLoS ONE, 6(1), e14592. 10.1371/journal.pone.0014592

Berger, D. J., German, D. W., John, C., Hart, R., Stephenson, T. R., & Avgar, T. (2022). Seeing is be- leaving: perception informs migratory decisions in Sierra Nevada Bighorn sheep (*Ovis canadensis sierrae*). Frontiers in Ecology and Evolution, 10.

Brønnvik, H., Nourani, E., Fiedler, W., & Flack, A. (2024). Experience reduces route selection for conspecifics by the collectively migrating white stork. Current Biology.

Bonte, D., Van Dyck, H., Bullock, J. M., Coulon, A., Delgado, M., Gibbs, M., Lehouck, V., Matthysen, E., Mustin, K., Saastamoinen, M., Schtickzelle, N., Stevens, V. M., Vandewoestijne, S., Baguette, M., Barton, K., Benton, T. G., Chaput-Bardy, A., Clobert, J., Dytham, C., … Travis, J. M. J. (2012). Costs of dispersal. Biological Reviews, 87(2), 290–312. 10.1111/j.1469-185X.2011.00201.x

Bracis, C., Bildstein, K. L., & Mueller, T. (2018). Revisitation analysis uncovers spatio-temporal patterns in animal movement data. Ecography, 41(11), 1801–1811. 10.1111/ecog.03618

Bracis, C., & Mueller, T. (2017). Memory, not just perception, plays an important role in terrestrial mammalian migration. Proceedings of the Royal Society B: Biological Sciences, 284(1855), 20170449. 10.1098/rspb.2017.0449

Calabrese, J. M., Fleming, C. H., & Gurarie, E. (2016). ctmm: An R package for analyzing animal relocation data as a continuous-time stochastic process. Methods in Ecology and Evolution, 7(9), 1124– 1132. 10.1111/2041-210X.12559

Dall, S. R., Giraldeau, L. A., Olsson, O., McNamara, J. M., & Stephens, D. W. (2005). Information and its use by animals in evolutionary ecology. Trends in ecology & evolution, 20(4), 187–193.

Dalziel, B. D., Morales, J. M., & Fryxell, J. M. (2008). Fitting probability distributions to animal movement trajectories: using artificial neural networks to link distance, resources, and memory. The American Naturalist, 172(2), 248–258.

Duchesne, T., Fortin, D., & Rivest, L.-P. (2015). Equivalence between Step Selection Functions and Biased Correlated Random Walks for Statistical Inference on Animal Movement. PLOS ONE, 10(4), e0122947. 10.1371/journal.pone.0122947

Darley-Hill, S., & Johnson, W. C. (1981). Acorn dispersal by the blue jay (*Cyanocitta cristata*). Oecologia, 50(2), 231–232. 10.1007/BF00348043

del Mar Delgado, M., Miranda, M., Alvarez, S. J., Gurarie, E., Fagan, W. F., Penteriani, V., di Virgilio, A., & Morales, J. M. (2018). The importance of individual variation in the dynamics of animal collective movements. Philosophical Transactions of the Royal Society B: Biological Sciences, 373(1746), 20170008. 10.1098/rstb.2017.0008

Doherty, T. S., & Driscoll, D. A. (2018). Coupling movement and landscape ecology for animal conservation in production landscapes. Proceedings of the Royal Society B: Biological Sciences, 285(1870), 20172272. 10.1098/rspb.2017.2272

Dominoni, D. M., Halfwerk, W., Baird, E., Buxton, R. T., Fernández-Juricic, E., Fristrup, K. M., … & Barber, J. R. (2020). Why conservation biology can benefit from sensory ecology. Nature Ecology & Evolution, 4(4), 502–511.

Dukas, R. (Ed.). (1998). Cognitive ecology: the evolutionary ecology of information processing and decision making. University of Chicago Press.

Dusenbery, D. B. (1992). Sensory ecology: how organisms acquire and respond to information. (1^st^ ed.). W. H. Freeman.

Edwards, M. A., & Derocher, A. E. (2015). Mating-related behaviour of grizzly bears inhabiting marginal habitat at the periphery of their North American range. Behavioural Processes, 111, 75–83.

Eisaguirre, J. M., Williams, P. J., & Hooten, M. B. (2024). Rayleigh step-selection functions and connections to continuous-time mechanistic movement models. Movement Ecology, 12(1), 14. 10.1186/s40462-023-00442-w

Ellison, N., Hatchwell, B.J., Biddiscombe, S.J., Napper, C.J. & Potts, J.R. (2020). Mechanistic home range analysis reveals drivers of space use patterns for a non-territorial passerine. J. Anim. Ecol., 89, 2763–2776.

Fagan, W. F., Lewis, M. A., Auger-Méthé, M., Avgar, T., Benhamou, S., Breed, G., LaDage, L., Schlägel, U. E., Tang, W., Papastamatiou, Y. P., Forester, J., & Mueller, T. (2013). Spatial memory and animal movement. Ecology Letters, 16(10), 1316–1329. 10.1111/ele.12165

Fagan, W. F., McBride, F., & Koralov, L. (2023). Reinforced diffusions as models of memory-mediated animal movement. Journal of the Royal Society Interface, 20(200), 20220700.

Fieberg, J., Signer, J., Smith, B., & Avgar, T. (2021). A ‘How to’ guide for interpreting parameters in habitat-selection analyses. Journal of Animal Ecology, 90(5), 1027–1043. 10.1111/1365-2656.13441

Fleming, C. H., Fagan, W. F., Mueller, T., Olson, K. A., Leimgruber, P., & Calabrese, J. M. (2015). Rigorous home range estimation with movement data: a new autocorrelated kernel density estimator. Ecology, 96(5), 1182–1188.

Finnegan, S. P., Svoboda, N. J., Fowler, N. L., Schooler, S. L., & Belant, J. L. (2021). Variable intraspecific space use supports optimality in an apex predator. Scientific Reports, 11(1), 21115. 10.1038/s41598-021-00667-y

Fleming, C. H., Fagan, W. F., Mueller, T., Olson, K. A., Leimgruber, P., & Calabrese, J. M. (2015). Rigorous home range estimation with movement data: A new autocorrelated kernel density estimator. Ecology, 96(5), 1182–1188. 10.1890/14-2010.1

Fortin, D., Beyer, H. L., Boyce, M. S., Smith, D. W., Duchesne, T., & Mao, J. S. (2005). Wolves influence elk movements: behavior shapes a trophic cascade in Yellowstone National Park. Ecology, 86(5), 1320–1330.

Forrest, S. W., Pagendam, D., Bode, M., Drovandi, C., Potts, J. R., Perry, J., … & Hoskins, A. J. (2024). Simulating animal movement trajectories from temporally dynamic step selection functions. bioRxiv, 2024-03.

García-Rodríguez, A., Selva, N., Zwijacz-Kozica, T., Albrecht, J., Lionnet, C., Rioux, D., Taberlet, P., & De Barba, M. (2021). The bear-berry connection: Ecological and management implications of brown bears’ food habits in a highly touristic protected area. Biological Conservation, 264, 109376. 10.1016/j.biocon.2021.109376

Gautestad, A. O., Loe, L. E., & Mysterud, A. (2013). Inferring spatial memory and spatiotemporal scaling from GPS data: comparing red deer Cervus elaphus movements with simulation models. Journal of animal Ecology, 82(3), 572–586.

Gehr, B., Bonnot, N., Heurich, M., Cagnacci, F., Ciuti, S., Hewison, A.J.M., et al. (2020). Stay home, stay safe - site familiarity reduces predation risk in a large herbivore in two contrasting study sites. J. Anim. Ecol., 89, 1329–1339

Gurarie, E., & Avgar, T. (2024). Cognitive movement ecology. Frontiers in Ecology and Evolution, 12, 1360427.

Gurarie, E., Bracis, C., Brilliantova, A., Kojola, I., Suutarinen, J., Ovaskainen, O., Potluri, S., & Fagan, W. F. (2022). Spatial memory drives foraging strategies of wolves, but in highly individual ways. Frontiers in Ecology and Evolution, 10. https://www.frontiersin.org/article/10.3389/fevo.2022.768478

Hayes, M. A. (2015). Dispersal and population genetic structure in two flyways of sandhill cranes *(Grus canadensis)*. The University of Wisconsin-Madison.

Heathcote, R. J., Whiteside, M. A., Beardsworth, C. E., Van Horik, J. O., Laker, P. R., Toledo, S., … & Madden, J. R. (2023). Spatial memory predicts home range size and predation risk in pheasants. Nature Ecology & Evolution, 7(3), 461–471.

Hofmann, D. D., Cozzi, G., McNutt, J.W., Ozgul, A., & Behr, D. M. (2023). A three-step approach for assessing landscape connectivity via simulated dispersal: African wild dog case study. Landscape Ecology, 38 (4), 981–998. 10.1007/s10980-023-01602-4

Hooten, M. B., Scharf, H. R., & Morales, J. M. (2019). Running on empty: Recharge dynamics from animal movement data. Ecology Letters, 22(2), 377–389. 10.1111/ele.13198

Janson, C. H., & Byrne, R. (2007). What wild primates know about resources: Opening up the black box. Animal Cognition, 10(3), 357–367. 10.1007/s10071-007-0080-9

Joly, K., Gurarie, E., Sorum, M. S., Kaczensky, P., Cameron, M. D., Jakes, A. F., Borg, B. L., Nandintsetseg, D., Hopcraft, J. G. C., Buuveibaatar, B., Jones, P. F., Mueller, T., Walzer, C., Olson, K. A., Payne, J. C., Yadamsuren, A., & Hebblewhite, M. (2019). Longest terrestrial migrations and movements around the world. Scientific Reports, 9(1), 15333. 10.1038/s41598-019-51884-5

Jordan, L. A., & Ryan, M. J. (2015). The sensory ecology of adaptive landscapes. Biology letters, 11(5), 20141054.

Kashetsky, T., Avgar, T., & Dukas, R. (2021). The cognitive ecology of animal movement: Evidence from birds and mammals. Frontiers in Ecology and Evolution, 9, 724887. 10.3389/fevo.2021.724887

Klappstein, N., Michelot, T., Fieberg, J., Pedersen, E., Field, C., & Flemming, J. M. (2024). Step selection functions with non-linear and random effects. Methods in Ecology and Evolution.

Lewis, M. A., Fagan, W. F., Auger-Méthé, M., Frair, J., Fryxell, J. M., Gros, C., Gurarie, E., Healy, S. D., & Merkle, J. A. (2021). Learning and animal movement. Frontiers in Ecology and Evolution, 9, 681704. 10.3389/fevo.2021.681704

Lima, S. L., & Zollner, P. A. (1996). Towards a behavioral ecology of ecological landscapes. Trends in Ecology & Evolution, 11(3), 131–135.

MacHutchon, A. G., & Wellwood, D. W. (2003). Grizzly Bear Food Habits in the Northern Yukon, Canada.

Matthiopoulos, J., J. Fieberg, and G. Aarts. (2023). Species-Habitat Associations: Spatial data, predictive models, and ecological insights, 2nd Edition. University of Minnesota Libraries Publishing. Retrieved from the University of Minnesota Digital Conservancy, https://hdl.handle.net/11299/217469.

Matthysen, E. (2005). Density-dependent dispersal in birds and mammals. Ecography, 28(3), 403–416. 10.1111/j.0906-7590.2005.04073.x

McLoughlin, C. (2002). Learner support in distance and networked learning environments: Ten dimensions for successful design. Distance Education, 23(2), 149–162. 10.1080/0158791022000009178

Merkle, J.A., Fortin, D. & Morales, J.M. (2014). A memory-based foraging tactic reveals an adaptive mechanism for restricted space use. Ecol. Lett., 17, 924–931.

Merkle, J. A., Potts, J. R., & Fortin, D. (2017). Energy benefits and emergent space use patterns of an empirically parameterized model of memory-based patch selection. Oikos, 126(2). 10.1111/oik.03356

Merkle, J. A., Sawyer, H., Monteith, K. L., Dwinnell, S. P. H., Fralick, G. L., & Kauffman, M. J. (2019). Spatial memory shapes migration and its benefits: Evidence from a large herbivore. Ecology Letters, 22(11), 1797–1805. 10.1111/ele.13362

Michelot, T., Klappstein, N. J., Potts, J. R., & Fieberg, J. (2024). Understanding step selection analysis through numerical integration. Methods in Ecology and Evolution, 15(1), 24–35.

Miles, D., Stedman, M., & Heald, A. (2020). Living with COVID-19: balancing costs against benefits in the face of the virus. National Institute Economic Review, 253, R60–R76.

Morrison, T. A., Merkle, J. A., Hopcraft, J. G. C., Aikens, E. O., Beck, J. L., Boone, R. B., Courtemanch, B., Dwinnell, S. P., Fairbanks, W. S., Griffith, B., Middleton, A. D., Monteith, K. L., Oates, B., Riotte-Lambert, L., Sawyer, H., Smith, K. T., Stabach, J. A., Taylor, K. L., & Kauffman, M. J. (2021). Drivers of site fidelity in ungulates. Journal of Animal Ecology, 90(4), 955–966. 10.1111/1365-2656.13425

Nabe-Nielsen, J., Tougaard, J., Teilmann, J., Lucke, K. & Forchhammer, M.C. (2013). How a simple adaptive foraging strategy can lead to emergent home ranges and increased food intake. Oikos, 122, 1307–1316.

Nathan, R., Getz, W. M., Revilla, E., Holyoak, M., Kadmon, R., Saltz, D., & Smouse, P. E. (2008). A movement ecology paradigm for unifying organismal movement research. Proceedings of the National Academy of Sciences, 105(49), 19052–19059. 10.1073/pnas.0800375105

Nathan, R., Monk, C. T., Arlinghaus, R., Adam, T., Alós, J., Assaf, M., Baktoft, H., Beardsworth, C. E., Bertram, M. G., Bijleveld, A. I., Brodin, T., Brooks, J. L., Campos-Candela, A., Cooke, S. J., Gjelland, K. Ø., Gupte, P. R., Harel, R., Hellström, G., Jeltsch, F., … Jarić, I. (2022). Big-data approaches lead to an increased understanding of the ecology of animal movement. Science, 375(6582), eabg1780. 10.1126/science.abg1780

Nesbitt, S. A. (1992). First reproductive success and individual productivity in Sandhill Cranes. The Journal of Wildlife Management, 56(3), 573. 10.2307/3808874

Northrup, J. M., Anderson Jr, C. R., & Wittemyer, G. (2016). Environmental dynamics and anthropogenic development alter philopatry and space-use in a North American cervid. Diversity and Distributions, 22(5), 547–557. 10.1111/ddi.12417

Oliveira-Santos, L. G. R., Forester, J. D., Piovezan, U., Tomas, W. M., & Fernandez, F. A. S. (2016). Incorporating animal spatial memory in step selection functions. Journal of Animal Ecology, 85(2), 516– 524. 10.1111/1365-2656.12485

Osipova, L., Okello, M. M., Njumbi, S. J., Ngene, S., Western, D., Hayward, M. W., & Balkenhol, N. (2019). Using step-selection functions to model landscape connectivity for African elephants: accounting for variability across individuals and seasons. Animal Conservation, 22(1), 35–48.

Pasitschniak-Arts, M. (1993). Ursus arctos. Mammalian Species, 439, 1–10.

Picardi, S., Abrahms, B., Gelzer, E., Morrison, T. A., Verzuh, T., & Merkle, J. A. (2023). Defining null expectations for animal site fidelity. Ecology Letters, 26(1), 157–169. 10.1111/ele.14148

Piper, W. H. (2011). Making habitat selection more “familiar”: a review. Behavioral Ecology and Sociobiology, 65, 1329–1351.

Potts, J. R., & Börger, L. (2023). How to scale up from animal movement decisions to spatiotemporal patterns: An approach via step selection. Journal of Animal Ecology, 92(1), 16–29.

Potts, J. R., & Lewis, M. A. (2016). How memory of direct animal interactions can lead to territorial pattern formation. Journal of The Royal Society Interface, 13(118), 20160059. 10.1098/rsif.2016.0059

Potts, J. R., & Schlägel, U. E. (2020). Parametrizing diffusion-taxis equations from animal movement trajectories using step selection analysis. Methods in Ecology and Evolution, 11(9), 1092–1105. 10.1111/2041-210X.13406

Ranc, N., Cagnacci, F., & Moorcroft, P. R. (2022). Memory drives the formation of animal home ranges: Evidence from a reintroduction. Ecology Letters, 25(4), 716–728. 10.1111/ele.13869

Ranc, N., Moorcroft, P. R., Ossi, F., & Cagnacci, F. (2021). Experimental evidence of memory-based foraging decisions in a large wild mammal. Proceedings of the National Academy of Sciences, 118(15), e2014856118. 10.1073/pnas.2014856118

Rheault, H., Anderson, C. R., Bonar, M., Marrotte, R. R., Ross, T. R., Wittemyer, G., & Northrup, J. M. (2021). Some memories never fade: Inferring multi-scale memory effects on habitat selection of a migratory ungulate using step-selection functions. Frontiers in Ecology and Evolution, 9. https://www.frontiersin.org/article/10.3389/fevo.2021.702818

Riotte-Lambert, L., Benhamou, S., & Chamaillé-Jammes, S. (2015). How memory-based movement leads to nonterritorial spatial segregation. The American Naturalist, 185(4), E103–E116. 10.1086/680009

Sawyer, H., LeBeau, C. W., McDonald, T. L., Xu, W., & Middleton, A. D. (2019). All routes are not created equal: An ungulate’s choice of migration route can influence its survival. Journal of Applied Ecology, 1365-2664.13445. 10.1111/1365-2664.13445

Sawyer, H., Middleton, A. D., Hayes, M. M., Kauffman, M. J., & Monteith, K. L. (2016). The extra mile: Ungulate migration distance alters the use of seasonal range and exposure to anthropogenic risk. Ecosphere, 7(10), e01534.

Schlägel, U. E., & Lewis, M. A. (2014). Detecting effects of spatial memory and dynamic information on animal movement decisions. Methods in Ecology and Evolution, 5(11), 1236–1246. 10.1111/2041-210X.12284

Schlägel, U. E., Merrill, E. H., & Lewis, M. A. (2017). Territory surveillance and prey management: Wolves keep track of space and time. Ecology and Evolution, 7(20), 8388–8405. 10.1002/ece3.3176

Selva, N., Teitelbaum, C. S., Sergiel, A., Zwijacz-Kozica, T., Zięba, F., Bojarska, K., & Mueller, T. (2017). Supplementary ungulate feeding affects movement behavior of brown bears. Basic and Applied Ecology, 24, 68–76. 10.1016/j.baae.2017.09.007

Signer, J., Fieberg, J., & Avgar, T. (2019). Animal movement tools (amt): R package for managing tracking data and conducting habitat selection analyses. Ecology and Evolution, 9(2), 880–890. 10.1002/ece3.4823

Sergio, F., Tavecchia, G., Blas, J., Lopez, L., Tanferna, A., & Hiraldo, F. (2011). Variation in age-structured vital rates of a long-lived raptor: implications for population growth. Basic and Applied Ecology, 12(2), 107–115.

Signer, J., Fieberg, J., Reineking, B., Schlaegel, U., Smith, B., Balkenhol, N., & Avgar, T. (2024). Simulating animal space use from fitted integrated Step-Selection Functions (iSSF). Methods in Ecology and Evolution, 15(1), 43–50.

Spencer, W. D. (2012). Home ranges and the value of spatial information. Journal of Mammalogy, 93(4), 929–947. 10.1644/12-MAMM-S-061.1

Subalusky, A. L., Dutton, C. L., Rosi, E. J., & Post, D. M. (2017). Annual mass drownings of the Serengeti wildebeest migration influence nutrient cycling and storage in the Mara River. Proceedings of the National Academy of Sciences, 114(29), 7647–7652. 10.1073/pnas.1614778114

Tacha, T. C., Haley, D. E., & Vohs, P. A. (1989). Age of sexual maturity of Sandhill Cranes from Mid-Continental North America. The Journal of Wildlife Management, 53(1), 43. 10.2307/3801303

Tello-Ramos, M. C., Branch, C. L., Kozlovsky, D. Y., Pitera, A. M., & Pravosudov, V. V. (2019). Spatial memory and cognitive flexibility trade-offs: To be or not to be flexible, that is the question. Animal Behaviour, 147, 129–136. 10.1016/j.anbehav.2018.02.019

Thompson, P. R., Derocher, A. E., Edwards, M. A., & Lewis, M. A. (2022). Detecting seasonal episodic-like spatio-temporal memory patterns using animal movement modelling. Methods in Ecology and Evolution, 13(1), 105–120. 10.1111/2041-210X.13743

Thompson, P. R., Lewis, M. A., Edwards, M. A., & Derocher, A. E. (2022). Time-dependent memory and individual variation in Arctic brown bears (*Ursus arctos*). Movement Ecology, 10(1), 18. 10.1186/s40462-022-00319-4

Thurfjell, H., Ciuti, S., & Boyce, M. S. (2014). Applications of step-selection functions in ecology and conservation. Movement Ecology, 2(1), 4. 10.1186/2051-3933-2-4

Trapanese, C., Meunier, H., & Masi, S. (2019). What, where and when: Spatial foraging decisions in primates: Spatial foraging decisions in primates. Biological Reviews, 94(2), 483–502. 10.1111/brv.12462

Tulving, E., & Craik, F. I. M. (2000). The Oxford Handbook of Memory. Oxford University Press.

Van Moorter, B., Visscher, D., Benhamou, S., Börger, L., Boyce, M. S., & Gaillard, J.-M. (2009). Memory keeps you at home: A mechanistic model for home range emergence. Oikos, 118(5), 641–652. 10.1111/j.1600-0706.2008.17003.x

Van Moorter, B., Visscher, D., Herfindal, I., Basille, M., & Mysterud, A. (2013). Inferring behavioural mechanisms in habitat selection studies getting the null-hypothesis right for functional and familiarity responses. Ecography, 36(3), 323–330.

Wang, H., & Salmaniw, Y. (2023). Open problems in PDE models for knowledge-based animal movement via nonlocal perception and cognitive mapping. Journal of Mathematical Biology, 86(5), 71. 10.1007/s00285-023-01905-9

Weise, F. J., Stratford, K. J., & R. J. van Vuuren. (2014). Financial costs of large carnivore translocations–accounting for conservation. PLoS One, 9(8), e105042.

Whittington, J., Hebblewhite, M., Baron, R. W., Ford, A. T., & Paczkowski, J. (2022). Towns and trails drive carnivore movement behaviour, resource selection, and connectivity. Movement Ecology, 10 (1), 17. 10.1186/s40462-022-00318-5

Wild, S., Alarcón-Nieto, G., Chimento, M., & Aplin, L. M. (2023). Manipulating actions: A selective two-option device for cognitive experiments in wild animals. Journal of Animal Ecology, 92(8), 1509–1519.

Wirsing, A. J., Quinn, T. P., Cunningham, C. J., Adams, J. R., Craig, A. D., & Waits, L. P. (2018). Alaskan brown bears (*Ursus arctos*) aggregate and display fidelity to foraging neighborhoods while preying on Pacific salmon along small streams. Ecology and Evolution, 8(17), 9048–9061. 10.1002/ece3.4431

Wolf, M., Frair, J., Merrill, E. & Turchin, P. (2009). The attraction of the known: The importance of spatial familiarity in habitat selection in wapiti *Cervus elaphu*s. Ecography, 32, 401–410.

Wolfson, D. W. (2018). Migratory ecology and movement patterns of mid-continent and eastern sandhill cranes. University of Minnesota.

Wolfson, D. W., Fieberg, J. R., & Andersen, D. E. (2020). Juvenile Sandhill Cranes exhibit wider ranging and more exploratory movements than adults during the breeding season. Ibis, 162(2), 556–562. 10.1111/ibi.12786

